# Expression reduction of biallelically transcribed X-linked genes during the human female preimplantation development

**DOI:** 10.1101/682286

**Authors:** Björn Reinius, Rickard Sandberg

## Abstract

Our previous single-cell RNA-seq data from human preimplantation embryos showed that female X-chromosome mRNA levels become partly dose compensated during the timespan between zygotic genome activation (ZGA) and implantation. At the same time, *XIST* RNA is expressed from, and forms clouds in proximity to, both X-chromosome copies and biallelic expression of other X-linked genes persists. We proposed that X-chromosome transcription is transiently lowered on both alleles before X-chromosome inactivation (XCI) takes place. This notion was recently challenged in a reanalysis performed by Moreira de Mello *et al*, claiming to provide evidence against biallelic expression dampening and that instead proper XCI was responsible for the observed dosage compensation. Here we have addressed this reanalysis and highlighted methodological issues, and we conclude a current lack of evidence against biallelic X-chromosome dampening.

## Introduction

We earlier provided a data resource containing 1,529 single-cell RNA-seq libraries spanning embryonic day (E) 3-7 of the human preimplantation development^1^. Analyzing these data, we uncovered for the first time that RNA levels of X-chromosome expression become partly dose compensated during the human female preimplantation development E4-E7, following the major wave of ZGA occurring E3-E4. Surprisingly, this dosage-compensation event took place even though key features of proper XCI were absent, since X-linked genes were biallelically deteted and *XIST* RNA was expressed from, and clouded, both X-chromosomes in most female E7 cells, as shown by our single-cell RNA-seq and RNA-FISH data. Interestingly, biallelic clouds of *XIST* simultaneous with biallelic primary transcript foci for unsilenced X-linked genes were reported also in an earlier RNA-FISH-based study^2^ on late human female preimplantation embryos, together with the lack of heterochromatic Barr bodies in female nuclei. Based on the single-cell RNA-seq and RNA-FISH results we proposed that the expression levels from both X chromosomes are transiently reduced in female preimplantation cells, preceding proper XCI through which near complete silencing of a single X chromosome transpires. An alternative, but perhaps more ambiguous, wording to describe this phenomenon would be that partial XCI initiates on both X-copies. We termed the transient biallelic expression reduction *“X-chromosome dampening”*. Transcriptionally dampened biallelic X-chromosome states were soon after reported in naïve human embryonic stem cells^3^, albeit at low cellular frequency under the available *in vitro* conditions. Intriguingly, Sousa *et al* recently reported^4^ that also mouse female embryonic cells at the onset of differentiation accumulate *Xist* on both X chromosomes, leading to reduced expression or partial silencing on both alleles before resorting to proper XCI.

Some time ago, Moreira de Mello *et al* published an article^5^ in *Scientific Reports* in which they claimed to have challenged the dampening hypothesis by a reanalysis of our human preimplantation single-cell RNA-seq data^1^ in conjunction with a dataset^6^ from Yan *et al*. This proposition was based on two main lines of evidences: (*1*) They reported a decrease in biallelic detection of X-linked transcripts containing single nucleotide polymorphisms (SNPs) in female cells E3-E7, albeit also a decrease in male cells, and claimed that this was indicative of XCI in females. (*2*) They reported that a set of X-linked genes, which they had classified as biallelically expressed, did *not* become dose-compensated during the preimplantation development. Altogether, they claimed that these data indicated proper XCI and refuted biallelic dampening. In this report, we reexamined Moreira de Mello *et al*’s analysis^5^ in the context of our single-cell RNA-seq data, and identified problems that raise questions on the validity of their conclusions.

## Results

### Changed rate of monoallelic or biallelic detection in single-cell RNA-seq libraries is not in itself indicative of XCI

The first argument against X-chromosome dampening provided by Moreira de Mello *et al* was a decrease in biallelic SNP detections, reported as a significant (p<0.001) negative correlation between rates of biallelic X-gene detection and embryonic time E3-E7 (Moreira de Mello 2017^5^ figure 1). Female cells had a Pearson correlation of −0.21 and male cells had a lower rate of biallelic detection and Pearson correlation of −0.25, and this was interpreted as a sign of XCI in females while the ongoing degradation of maternal RNA in males. These patterns are clearly very different from those obtained in our own allelic analyses of the same data. In males, we detected a drastic clearance of biallelic X-chromosome RNAs between E3 and E4 (convergence close to zero at E4) along with the activation of paternally derived genes (Petropoulos 2016^1^ figure 1D and S1H). We interpreted this as the completion of the major wave of ZGA and clearance of maternally derived RNA between E3-E4, in line with previous results^6^. In female cells, we detected a steady rate of biallelic X-chromosome expression throughout the preimplantation development, which did not differ markedly from that of autosomal chromosomes (Petropoulos 2016^1^ figure 6). It is not worthwhile to here discuss what analysis procedure (SNP-filtering, thresholds etc.) might be the more appropriate for the allele-specific analysis of single-cell RNA-seq data, since this issue is complex and involves a high degree of uncertainty given that the human preimplantation embryos were of unknown and variable haplotypes. Instead, we will directly demonstrate why a slight potential decrease in biallelic detection in single-cell RNA-seq libraries is in itself not *“indicative of XCI”*, as claimed by Moreira de Mello *et al*.

**Figure 1.**
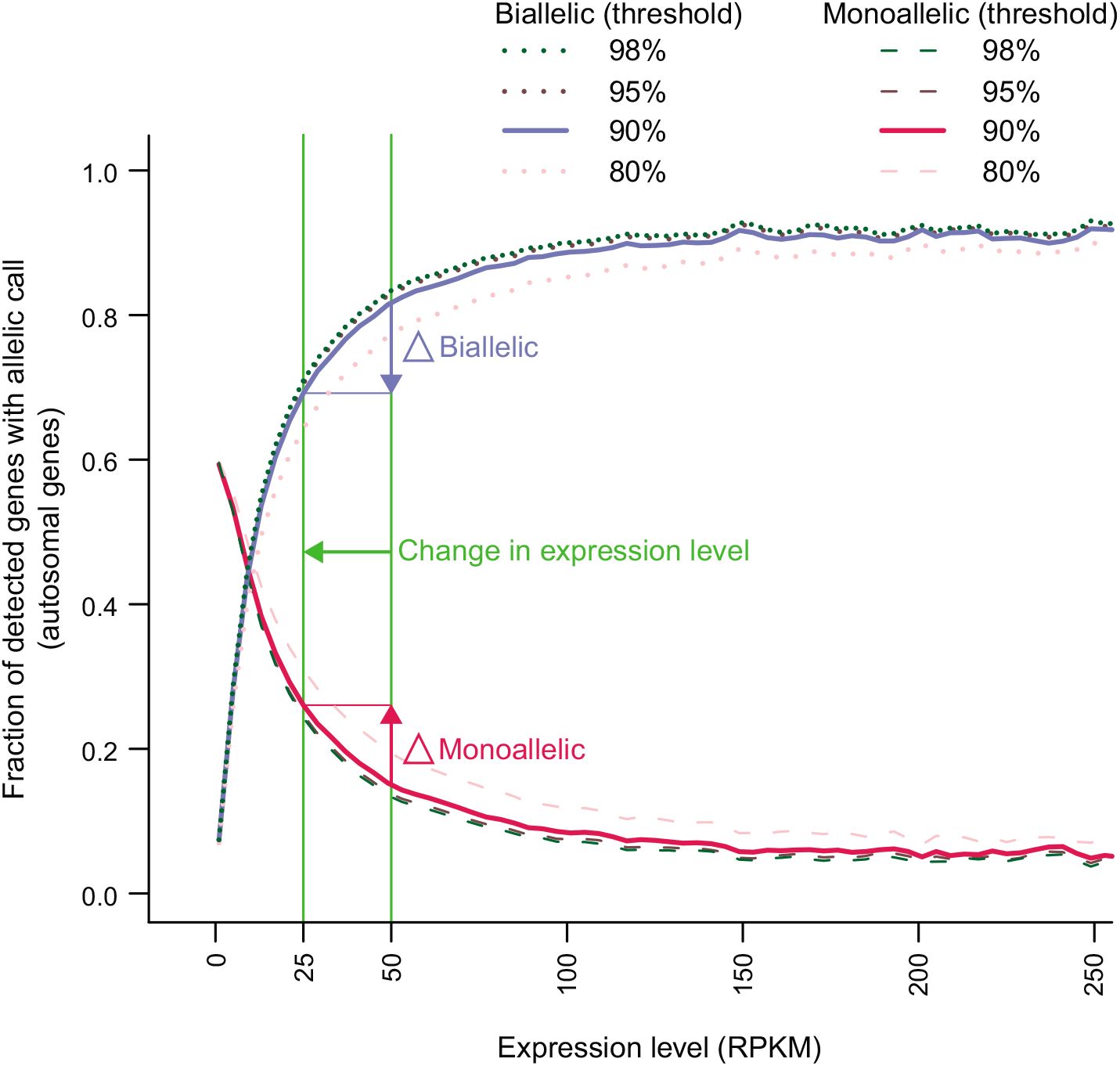
Probability of allelic calls in single-cell RNA-seq libraries. Fraction of genes (y-axis) with different allelic calls in single-cell RNA-seq libraries (CAST/EiJ × C57BL/6J mouse fibroblasts) as a function of gene expression level (x-axis). Genes were binning by their expression (RPKM) in single fibroblast cells sequenced using the Smart-seq2 method, and the fraction of each allelic call in expression-level bins was calculated. Thresholds correspond to the minimum allelic bias for calling expression “monoallelic”. Only autosomal (i.e. bialellically expressed) genes are included in the figure. High-expressed genes are generally detected as biallelic while low-expressed genes are more often monoallelically detected, due to technical dropout and transcriptional fluctuation^12^ (i.e. “bursting”). If the expression level of a gene, or subset of genes, is reduced (green arrow and vertical lines) the fraction of monoallelic detections in the single-cell RNA-seq library increases and the fraction biallelic detections decreases – even though the genes are still biallelically expressed in cells over time. The data and figure is adapted from Reinius et al 2016^9^.

It is now well understood that the probability of detecting biallelic expression of a gene in single-cell RNA-seq (including autosomal genes) is directly dependent on the gene’s overall expression level in the cell^7–11^. This can be understood on the one hand by a biological mechanism, as high-expressed genes have a higher probability to produce transcriptional bursts simultaneously from both alleles at any given time-window of sampling than low-expressed genes^12^. Additionally, RNA molecules present in low copy numbers are more prone to technical dropout during the single-cell library preparation and sequencing, which further exaggerates monoallelic observation. This dependence is demonstrated in **Fig. 1**, which shows the fraction biallelic and monoallelic detection in single-cell RNA-seq data as a function of average expression level for autosomal genes (data^9^ from hybrid [CAST/EiJ × C57BL/6J] mouse cells with well-characterized genomes). If the expression level of a gene, or a collection of genes, is shifted to lower level (**Fig. 1**, green arrow) the rate of monoallelic calls increases (**Fig. 1**, red arrow) and biallelic calls decreases (**Fig. 1**, blue arrow) – even though the genes are biallelically expressed in the cells over time. The premise in the analysis by Moreira de Mello *et al* was that X-chromosome gene expression levels are lowered between E4-E7 in female human preimplantation embryos^1^, either by biallelic dampening or by ongoing proper XCI. A potential decrease in biallelic X-gene detection during this time-window is thus by itself strictly insufficient to discern between the models. To indicate XCI using single-cell RNA-seq data it would instead be necessary to show a decrease in biallelic detection that is greater than what is expected from the reduction of expression levels alone. This might be approximated from autosomal expression-level distributions, but none of that were accounted for or even raised as a caveat in the analysis by Moreira de Mello *et al*. Altogether, the allelic expression analyses presented by Moreira de Mello *et al* are inconclusive and lack the proper controls. A similar issue in the analysis of rates of biallelic expression during the preimplantation development is present in a paper from Vallot *et al*^13^ (Vallot 2017^13^ figure 1).

Finally, we visited Supplementary Fig. S4 in Moreira de Mello *et al* and found that 7/22 autosomal chromosomes were reported to have a “significant” change in fraction biallelically called genes among female embryos during the preimplantation development (5 autosomes having positive and 2 having negative correlation; “Dataset #1”, Petropoulos data). In the same supplementary figure and for female embryos in the Yan dataset (“Dataset #2”) 12/22 autosomal chromosomes had “significant” negative change in fraction biallelically called genes. Because there is no chromosome-wide silencing of autosomes during the preimplantation development, these numbers serve as a measure of the false-positive rate in Moreira de Mello *et al*’s allelic analysis. This suggests a false discovery rate between ^~^9% (2/22) and ^~^55% (12/22) for a reported reduction in biallelic expression of a chromosome.

### Biallelically called X-chromosome genes are dose compensated

The second argument against X-chromosome dampening provided by Moreira de Mello *et al* was that genes which they had called as “biallelic” (thereby not subjected to XCI) did not become dose-compensated during the preimplantation development. We requested and obtained the lists of biallelically called genes from Moreira de Mello *et al*. (**Supplementary Table 1**, containing genes called as “biallelic” per embryonic day E3-E7). We then investigated in our single-cell expression data whether these “biallelic” genes tended to become down-regulated in female cells E4-E7 (i.e. following ZGA), which would be in line with dampening. Indeed, we found that this was the case, as reflected by a negative shift in the distribution of Spearman correlations between mean expression levels (Reads Per Kilobase of transcript per Million reads [RPKM]) and embryonic time exclusively for X-linked genes in females and not in males (p≤0.001, one-sided Wilcoxon test, **Fig. 2**). Note that it is not expected that all X-linked genes in females would have a negative Spearman correlation with embryonic time in this analysis, since many genes are up- or down-regulated as a function of embryonic development independent of any connection to dosage compensation. What we do expect in a scenario of dosage compensation is a negative shift in the distribution of Spearman correlation values, which is what we observed in female cells (**Fig. 2**).

**Figure 2.**
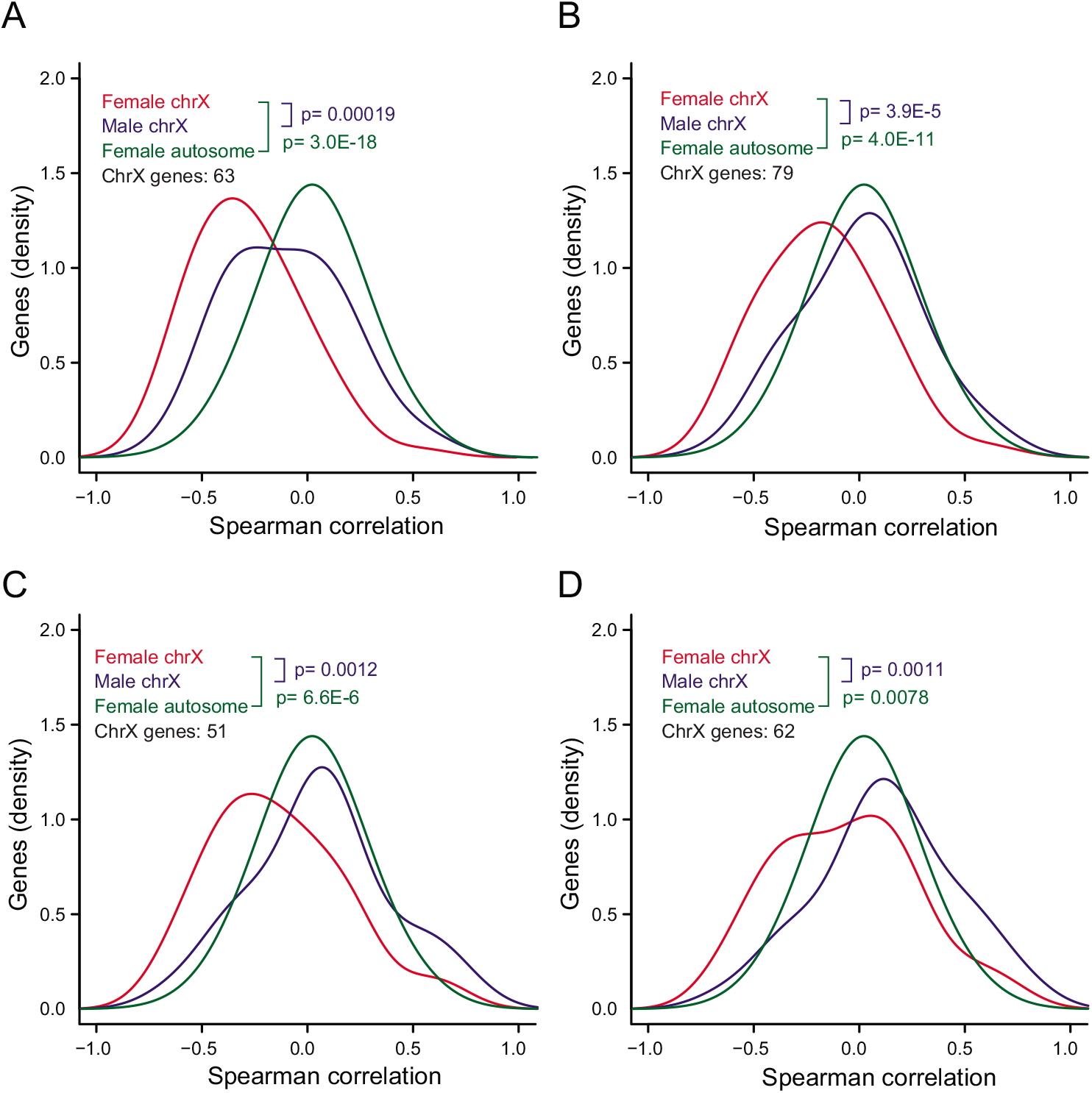
Expression trend of biallelic X-linked genes. Distribution (smoothed kernel density plot) of Spearman correlations between mean gene-expression levels and embryonic day (E4-E7) in female and male cells, for genes located on the X chromosome or autosomes. This is shown for genes called as biallelic in E4, E5, E6, and E7 in panel A, B, C, and D, respectively. P-values (female chrX vs male chrX in blue text, and female chrX vs female autosome in green text) according to one-sided Wilcoxon tests. Biallelic X-linked genes tended to have negative Spearman correlation.

We then investigated mean female-to-male relative expression levels for the “biallelic” genes, and in conflict with results of Moreira de Mello *et al* we again observed the pattern of dosage compensation E4-E7 (p≤1.8×10^−8^, two-sided Wilcoxon test, **Fig. 3**). Notably, female-to-male expression ratios have the benefit of being in part normalized for stage-specific gene-expression changes that are unrelated to dosage compensation, under assumption that the overall transcriptional differentiation program is similar in females and males during these stages.

**Figure 3.**
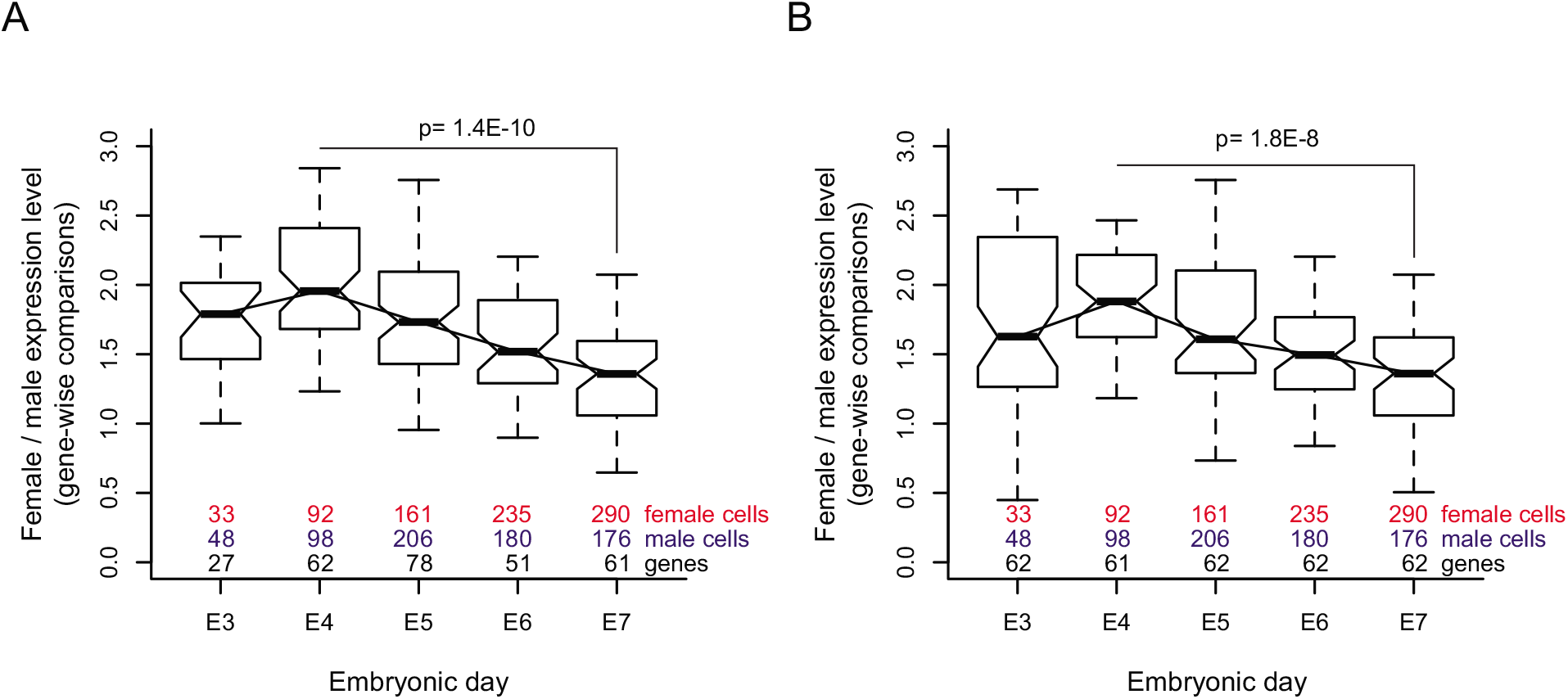
Female-to-male relative expression levels of biallelic X-linked genes. Boxplots of female-to-male expression-level ratios of biallelically called X chromosome genes, shown for genes called as biallelic at: **(A)** individual embryonic days, or **(B)** called as biallelic at E7. Genes with mean expression level above 1 RPKM per stage were included. P-values according to two-sided Wilcoxon tests. Boxes indicate the interquartile range, the belts indicate medians, and whiskers indicate extreme values at a maximum distance of the interquantile range.

Next, we checked whether the total (summed) expression of the “biallelic” gene set was reduced E4-E7. Moreira de Mello *et al* compared the total and median expression between *different* gene sets (genes called “biallelic” in different stages) over different embryonic time-points (Moreira de Mello 2017^5^ figure 3B-C). We, however, do not find this analysis sensible for detecting dosage compensation, since: (*1*) diverse genes are expressed at vastly different levels without any connection or relevance for dosage compensation, and (*2*) many genes change drastically (some beyond 100-fold^1^) in expression levels between E4-E7 as result of the developmental program rather than dosage compensation – which has an expected ≤2-fold effect. When comparing small sets of diverse genes, such as the current “biallelic” sets, it is therefore likely that dosage-compensation effects on sum or median expression become obfuscated by the mentioned factors if not accounted for. It is also important to appreciate that a handful of high-expressed genes may completely dominate the expression-level sums if an upper threshold is not imposed to remove outlier genes. Finally, Moreira de Mello *et al did* report a decreased total expression of “biallelic” genes, but then questioned whether this change was the result of a *decreased number* or *decreased expression levels* of “biallelic” genes with time – which we found to be a somewhat puzzling question given that they should know exactly the number of genes they included in each of their plots and calculations (Moreira de Mello 2017^5^ figure 3). The mentioned issues considered, we calculated the total expression of the fixed set of genes that were biallelically called at E7 over embryonic time (**Fig. 4**), reasoning that if these genes were not subjected to XCI at E7 they would also not be subjected to XCI at earlier stages. After the removal of high-expressed outlier genes (genes with expression levels beyond the 90th percentile – which dominated the total expression, **Fig. 4A-B**), we observed the signature of dosage compensation E4-E7 also in terms of their total expression output from the X-chromosome (p=2.8×10^−27^, two-sided Wilcoxon test, **Fig. 4C-D**). As expected, using the Transcripts Per Million reads (TPM) expression measure instead of RPKM provided very similar results (**Supplementary Fig. 1**).

**Figure 4.**
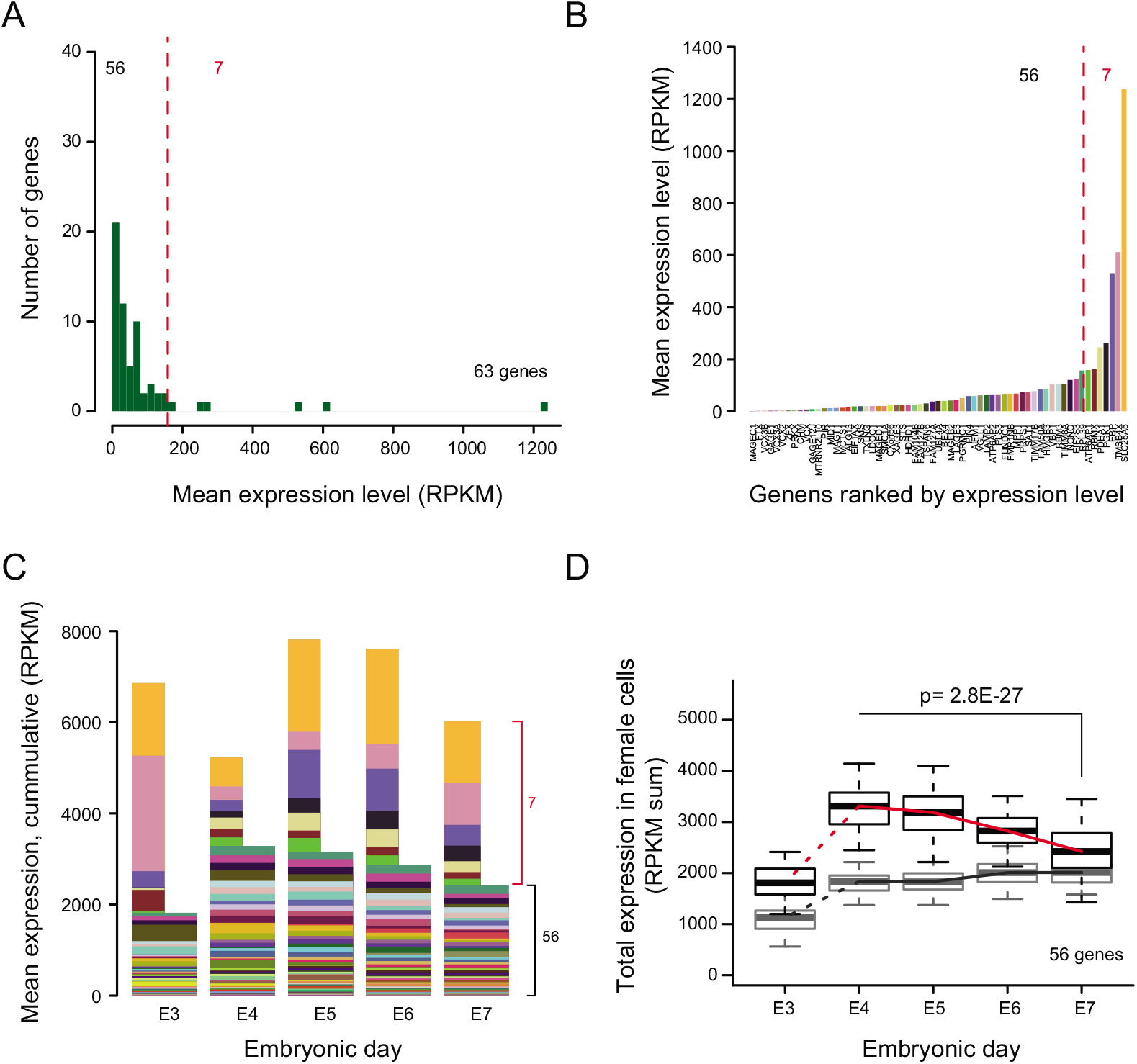
Total expression of biallelic X-linked genes. **(A)** Histogram of mean expression levels of genes called as “biallelic” at E7 (n=63) in female cells. The vertical red line indicates the 90th percentile. **(B)** Biallelic E7 genes ranked according to mean expression level. **(C)** Cumulative expression of biallelic E7 genes, demonstrating that a few high-expressed outlier genes (n=7, genes beyond 90th percentile, narrow bars) may dominate the total expression estimate and interpretation of expression trend. **(D)** Boxplots showing the total expression level (summed expression of 56 biallelic E7 genes; 90% of genes) in female cells at different stages of preimplantation development (x-axis); P-value for the comparison female E4:E7 according to a two-sided Wilcoxon test. Total expression in male cells is included as a gray boxplots.

Altogether, our reanalyzes showed that biallelically called X-chromosome genes tended to get dose compensated between E4-E7, as reflected by (*1*) a negative shift in the distribution of Spearman correlations between expression level and embryonic time; (*2*) decreasing female-to-male expression ratios; and (*3*) decreasing total expression.

It is possible that the divide in results between us^1^ and Moreira de Mello *et al*^5^ derive from differences in sequence-read mapping procedures and/or gene-expression-level calculations. Sample selection could be yet another reason, as we noticed that Moreira de Mello *et al* excluded about a third of all the single-cell libraries available in our resource. We therefore requested and obtained the processed gene-expression-level (TPM) data and lists of included samples from Moreira de Mello *et al*, and again performed the above described analyses. Interestingly, we found the signatures of dosage compensation in “biallelic” genes also in Moreira de Mello *et al*’s processed data set, including the reduction of female-to-male expression ratios (**Supplementary Fig. 2**).

## Conclusions

The presence of an initial biallelic X-chromosome dampening during the human female preimplantation development is an interesting and perhaps provoking thesis under some prevailing dogma in the XCI field, and as such needs to get challenged and tested^14^. However, we did not find the claims, nor the analysis, by Moreira de Mello *et al*^5^ convincing. We would like to emphasize that we do not dismiss the possibility of XCI initiating at some point, or in some fraction of cells, during the preimplantation development. Nevertheless, *“compelling evidence against X dampening in human preimplantation embryos”* as claimed^5^ by Moreira de Mello *et al* is currently lacking. Furthermore, biallelic dampening and XCI should not be considered mutually exclusive phenomena, but rather that biallelic expression reduction may precede proper XCI, as proposed in our original manuscript^1^ and recently reported to occur in some degree also in mouse embryonic cells^4^. It is known since long that expression of a translocated *Xist* gene induces cloud formation and expression reduction or silencing in *cis* even on autosomal chromosomes^15^.

We propose that dual X-chromosome dampening, which correlates with the appearance of biallelic *XIST* clouds in female cells, should be experimentally tested in human embryos for which the maternal and paternal X-chromosome haplotypes have been characterized. Such studies should preferably aim at the E4-E6 time-window, as the major wave of ZGA completes around E4 and proper XCI is likely to commence at some point around the time of implantation. Further studies on naïve human embryonic stem cells that express *XIST* from both alleles^3^ would also be valuable. If such analyses are performed using single-cell RNA-seq it is vital that the technical and biological stochastic features present in single-cell data are carefully considered.

## Methods

### Access to data

Calculations of expression levels (RPKMs) are described earlier^1, 16^. Expression-level data are available together with sample annotations at Sourceforge (https://sourceforge.net/projects/dampening-of-biallelic-genes/files/). Expression levels (TPM) calculated by Moreira de Mello *et al*, sample annotations, and lists of “biallelic” genes (**Supplementary Table 1**) were requested and obtained from Joana C. Moreira de Mello and Lygia V. Pereira, and are also available at Sourceforge.

### Spearman correlations between expression levels and embryonic time

We included biallelically called genes (**Supplementary Table 1**) with a mean expression level above 1 RPKM or 1 TPM (E4-E7) and calculated and plotted a smoothed kernel density (bandwidth= 0.15). We noted a negative shift in the distribution for female X-linked genes, and tested whether the median of female X-linked genes was significantly lower than that of female autosomal and male X-linked genes. P-values for differences in medians were calculated using a one-sided Wilcoxon tests. Using an expression-level cutoff of 5 RPKM instead of 1 RPKM provided similar results (**Supplementary Fig. 3A-D**).

### Female-to-male relative expression levels

We included biallelically called genes (**Supplementary Table 1**) with a mean expression level above 1 RPKM or 1 TPM (stage-wise for genes called “biallelic” per stage, and E3-E7 for fixed sets of biallelic genes) and calculated the gene-wise female/male ratios of mean expression levels. Ratios of finite value were included in the plots. P-values for differences in medians between E4 and E7 were calculated using a two-sided Wilcoxon test. Using an expression-level cutoff of 5 RPKM provided similar results (**Supplementary Fig. 3E-F**).

### Total expression

We calculated the mean expression level for each biallelically called E7 gene (**Supplementary Table 1**) and removed genes with mean RPKM or TPM beyond the 90^th^ percentile of expression levels (**Fig. 4A-C**) since such genes would otherwise dominate the total expression estimate. The sum of expression levels (RPKM) was then calculated in each cell for the remaining 90% of genes (**Fig. 4D**). Difference in summed expression between E4 and E7 for female cell was calculated using a two-sided Wilcoxon test.

## Code accessibility

Computational code used in this analysis is available at Sourceforge (https://sourceforge.net/projects/dampening-of-biallelic-genes/files/).

## Author contributions

B.R. performed the analyses, interpreted the results, drafted and edited the manuscript. R.S. interpreted the results and edited the manuscript.

## Supplementary Information

Supplementary Table 1 and Supplementary Figure 1-3.

## Acknowledgements

This work was supported by funding from the Ragnar Söderberg Foundation, the Swedish Research Council (2017-01723), and Åke Wiberg’s Foundation to BR; and the European Research Council (CoG 648842) to RS. This paper is under review in Scientific Reports (submitted July 10, 2018).

