## Supplementary Figures and Tables for "Expression reduction of biallelically transcribed X-linked genes during the human female preimplantation development"

**Contents:** Supplementary Figure 1-3, Supplementary Table 1

### Supplementary Figure 1

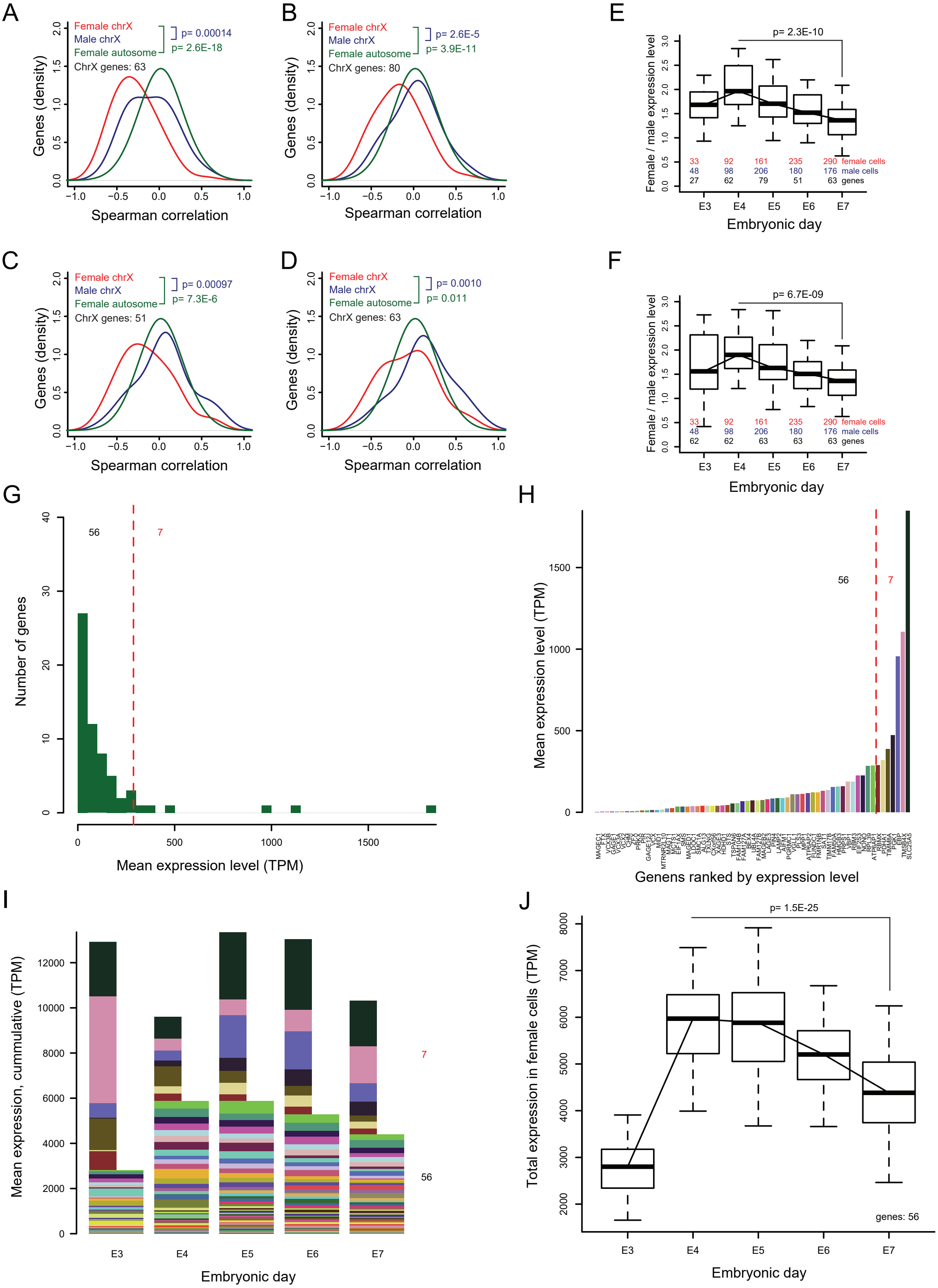


**Supplementary Figure 1.**
The same analysis as presented in main Figure 2-4, but using TPMs instead of RPKMs as expression-level measure. (A-D) As in main Figure 2. (E-F) As in main Figure 3. (G-J) as in main Figure 4.

### Supplementary Figure 2

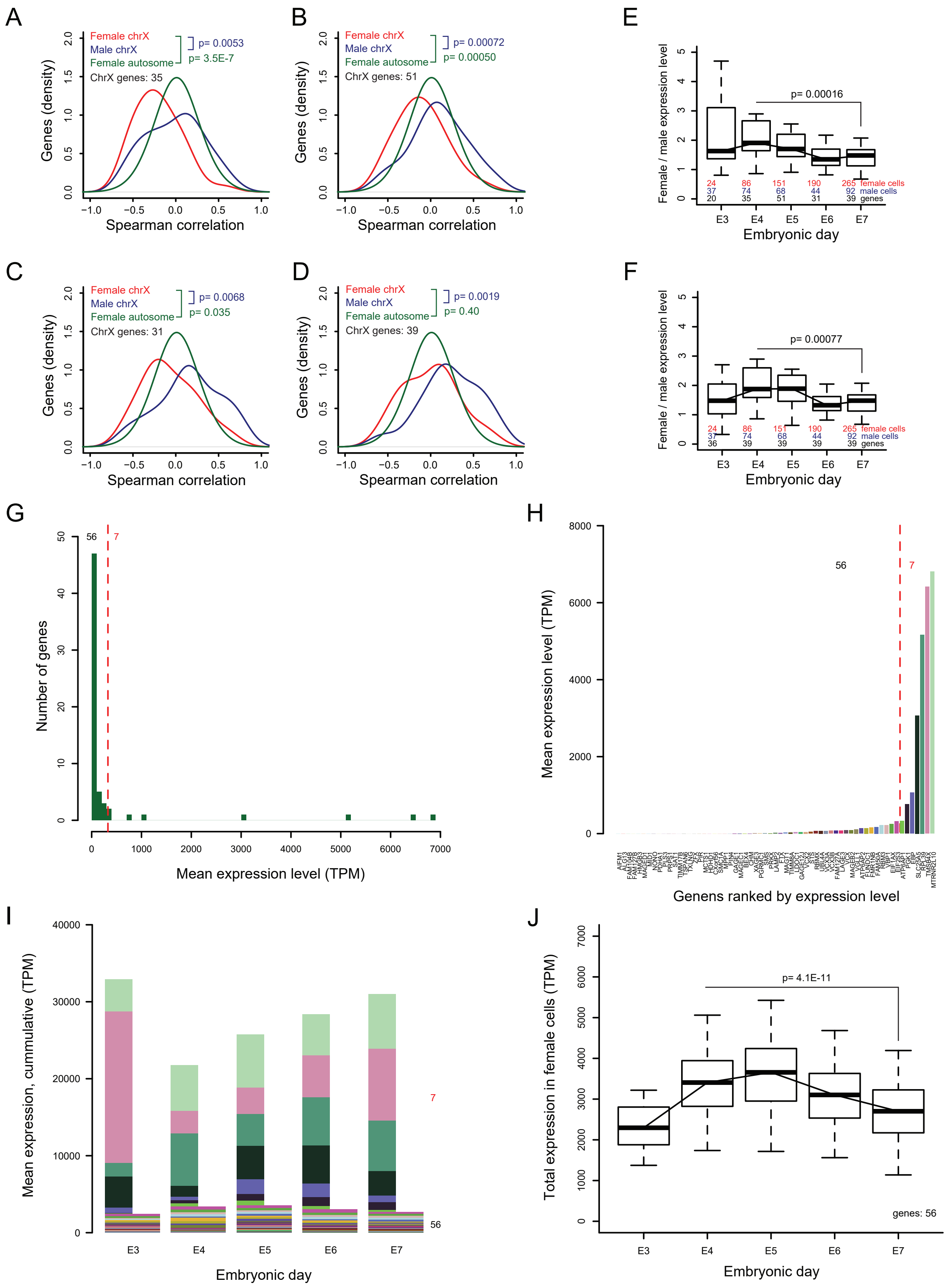


**Supplementary Figure 2.**
The same analysis as presented in main Figure 2-4, but using TPMs and sample lists obtained from Moreira de Mello *et al*. (A-D) As in main Figure 2. (E-F) As in main Figure 3. (G-J) as in main Figure 4.

### Supplementary Figure 3

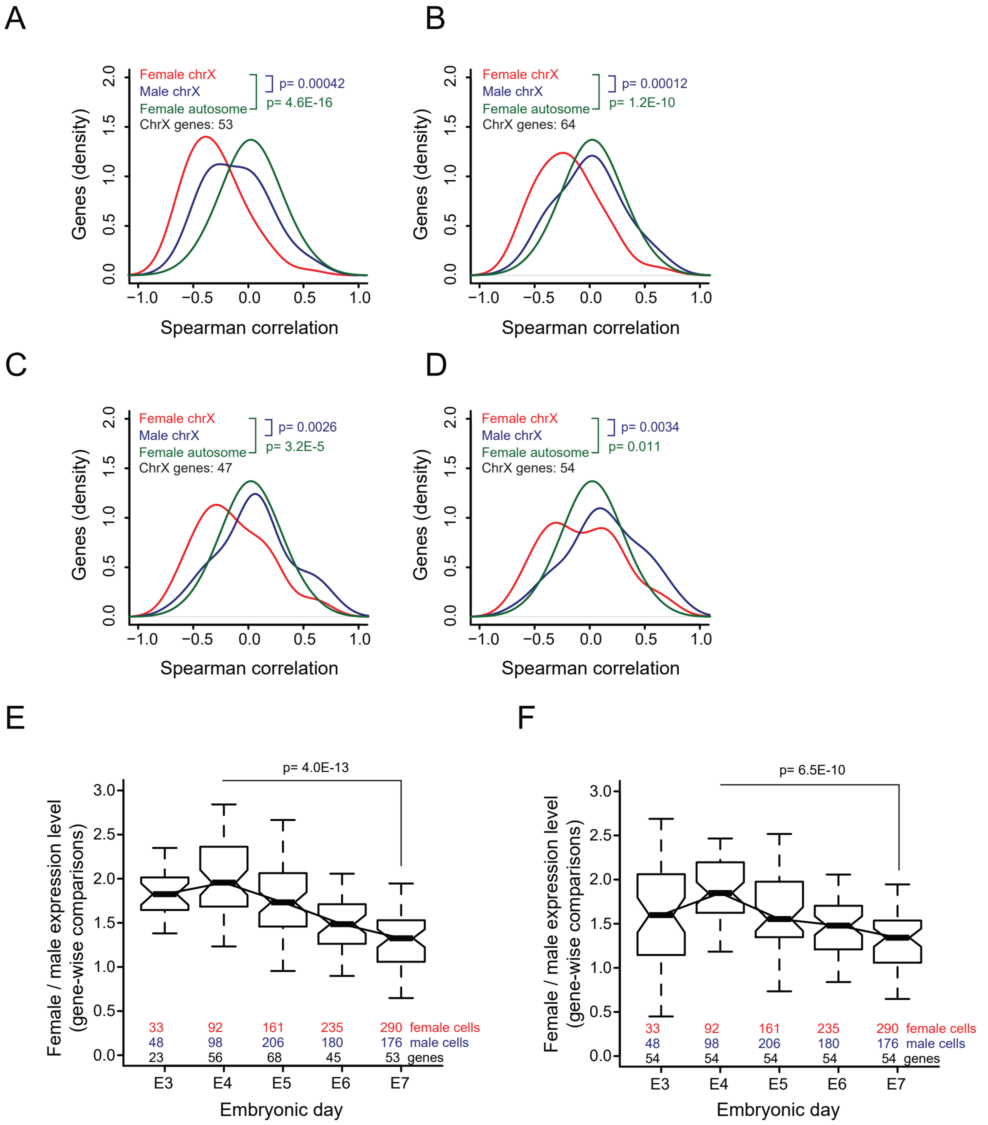


**Supplementary Figure 3.**
The same analysis as presented in main Figure 2-3, but including only genes with mean expression above RPKM 5. (A-D) as in main Figure 2. (E-F) as in main Figure 3.

**Supplementary Table 1**

**Genes called as biallelic at E3 from Moreira de Mello.**

| Chromosome | Gene symbol | RefSeq accession | in.E3.list | in.E4.list | in.E5.list | in.E6.list | in.E7.list |
|---|---|---|---|---|---|---|---|
| chrX | APOO | NR_026545 | yes | yes | yes | no | no |
| chrX | ATP6AP2 | NM_005765 | yes | no | no | no | yes |
| chrX | BMP15 | NM_005448 | yes | no | no | no | no |
| chrX | CHM | NM_000390 | yes | no | yes | no | yes |
| chrX | CSAG1 | NM_153478 | yes | yes | no | no | no |
| chrX | CSTF2 | NM_001325 | yes | no | no | no | no |
| chrX | CXorf56 | NM_001170569 | yes | yes | yes | no | yes |
| chrX | DKC1 | NM_001142463 | yes | yes | yes | yes | no |
| chrX | EBP | NM_006579 | yes | yes | yes | yes | yes |
| chrX | EIF2S3 | NM_001415 | yes | yes | yes | yes | yes |
| chrX | HCCS | NM_005333 | yes | yes | no | no | no |
| chrX | MAGEA4 | NM_002362 | yes | yes | yes | yes | no |
| chrX | MAGT1 | NM_032121 | yes | no | yes | no | yes |
| chrX | MOSPD1 | NM_019556 | yes | yes | no | no | no |
| chrX | MTRNR2L10 | NM_001190708 | yes | yes | yes | yes | yes |
| chrX | NDUFA1 | NM_004541 | yes | no | no | no | no |
| chrX | NONO | NM_001145410 | yes | yes | yes | yes | yes |
| chrX | PQBP1 | NM_001167992 | yes | yes | yes | no | no |
| chrX | RBM3 | NM_006743 | yes | yes | yes | yes | yes |
| chrX | RBMX | NM_001164803 | yes | yes | yes | yes | yes |
| chrX | RPL39 | NM_001000 | yes | yes | yes | yes | yes |
| chrX | SLC25A5 | NM_001152 | yes | yes | yes | yes | yes |
| chrX | TCEANC | NM_152634 | yes | no | no | no | no |
| chrX | TSIX | - | yes | yes | yes | yes | yes |
| chrX | VCX | NM_013452 | yes | yes | yes | yes | yes |
| chrX | VCX2 | NM_016378 | yes | yes | yes | yes | yes |
| chrX | VCX3A | NM_016379 | yes | yes | yes | yes | yes |
| chrX | VCX3B | NM_001001888 | yes | yes | yes | yes | yes |

**Genes called as biallelic at E4 from Moreira de Mello.**

| Chromosome | Gene symbol | RefSeq accession | in.E3.list | in.E4.list | in.E5.list | in.E6.list | in.E7.list |
|---|---|---|---|---|---|---|---|
| chrX | AIFM1 | NM_001130847 | no | yes | yes | yes | yes |
| chrX | APOO | NR_026545 | yes | yes | yes | no | no |
| chrX | ATP6AP1 | NM_001183 | no | yes | yes | yes | yes |
| chrX | BEX4 | NM_001080425 | no | yes | yes | yes | yes |
| chrX | BRWD3 | NM_153252 | no | yes | yes | no | no |
| chrX | CA5B | NM_007220 | no | yes | yes | no | no |
| chrX | CETN2 | NM_004344 | no | yes | no | no | no |
| chrX | CSAG1 | NM_153478 | yes | yes | no | no | no |
| chrX | CXorf38 | NM_144970 | no | yes | yes | no | no |
| chrX | CXorf56 | NM_001170569 | yes | yes | yes | no | yes |
| chrX | DKC1 | NM_001142463 | yes | yes | yes | yes | no |
| chrX | EBP | NM_006579 | yes | yes | yes | yes | yes |
| chrX | EIF1AX | NM_001412 | no | yes | yes | yes | yes |
| chrX | EIF2S3 | NM_001415 | yes | yes | yes | yes | yes |
| chrX | EMD | NM_000117 | no | yes | yes | no | no |
| chrX | FAM122C | NM_001170784 | no | yes | no | no | no |
| chrX | FMR1NB | NM_152578 | no | yes | yes | yes | yes |
| chrX | GAGE1 | NR_102272 | no | yes | yes | yes | yes |
| chrX | GAGE12J | NM_001098406 | no | yes | yes | yes | yes |
| chrX | GDPD2 | NM_001171192 | no | yes | yes | yes | no |
| chrX | HCCS | NM_005333 | yes | yes | no | no | no |
| chrX | HDAC8 | NM_001166448 | no | yes | no | no | no |
| chrX | HMGB3 | NM_005342 | no | yes | yes | yes | yes |
| chrX | KIAA1210 | - | no | yes | no | no | no |
| chrX | LAGE3 | NM_006014 | no | yes | yes | yes | yes |
| chrX | MAGEA1 | NM_004988 | no | yes | no | no | no |
| chrX | MAGEA4 | NM_002362 | yes | yes | yes | yes | no |
| chrX | MAGEB2 | NM_002364 | no | yes | yes | yes | yes |

| | | | | | | | |
|---|---|---|---|---|---|---|---|
| chrX | MAP7D3 | NM_024597 | no | yes | yes | no | no |
| chrX | MMGT1 | NM_173470 | no | yes | no | yes | no |
| chrX | MOSPD1 | NM_019556 | yes | yes | no | no | no |
| chrX | MTRNR2L10 | NM_001190708 | yes | yes | yes | yes | yes |
| chrX | NAP1L6 | - | no | yes | no | no | no |
| chrX | NONO | NM_001145410 | yes | yes | yes | yes | yes |
| chrX | NXT2 | NM_018698 | no | yes | no | no | no |
| chrX | PAGE2B | NM_001015038 | no | yes | no | no | no |
| chrX | PHF8 | NM_001184896 | no | yes | no | no | no |
| chrX | PIN4 | NM_001170747 | no | yes | no | no | yes |
| chrX | PLS3 | NM_005032 | no | yes | yes | yes | yes |
| chrX | PNMA5 | NM_052926 | no | yes | no | no | no |
| chrX | PQBP1 | NM_001167992 | yes | yes | yes | no | no |
| chrX | RAB9A | NM_001195328 | no | yes | yes | no | no |
| chrX | RBM3 | NM_006743 | yes | yes | yes | yes | yes |
| chrX | RBMX | NM_001164803 | yes | yes | yes | yes | yes |
| chrX | RPL39 | NM_001000 | yes | yes | yes | yes | yes |
| chrX | SAT1 | NM_002970 | no | yes | yes | yes | yes |
| chrX | SLC25A5 | NM_001152 | yes | yes | yes | yes | yes |
| chrX | SSX3 | NM_021014 | no | yes | no | no | no |
| chrX | TIMM17B | NM_005834 | no | yes | yes | yes | yes |
| chrX | TIMM8A | NM_001145951 | no | yes | yes | yes | yes |
| chrX | TKTL1 | NM_001145934 | no | yes | yes | no | no |
| chrX | TMEM185A | NM_001174092 | no | yes | no | no | no |
| chrX | TMSB4X | NM_021109 | no | yes | yes | yes | yes |
| chrX | TSIX | - | yes | yes | yes | yes | yes |
| chrX | TSPAN6 | NM_001278743 | no | yes | yes | yes | yes |
| chrX | TSR2 | NM_058163 | no | yes | no | no | no |
| chrX | UBA1 | NM_153280 | no | yes | no | yes | no |
| chrX | VBP1 | NM_003372 | no | yes | yes | yes | yes |
| chrX | VCX | NM_013452 | yes | yes | yes | yes | yes |
| chrX | VCX2 | NM_016378 | yes | yes | yes | yes | yes |
| chrX | VCX3A | NM_016379 | yes | yes | yes | yes | yes |
| chrX | VCX3B | NM_001001888 | yes | yes | yes | yes | yes |
| chrX | XAGE3 | NM_133179 | no | yes | yes | yes | yes |
| chrX | XIAP | NM_001167 | no | yes | no | no | no |
| chrX | ZDHHC9 | NM_016032 | no | yes | no | no | no |
| chrX | ZFX | NM_003410 | no | yes | no | no | yes |

**Genes called as biallelic at E5 from Moreira de Mello.**

| Chromosome | Gene symbol | RefSeq accession | in.E3.list | in.E4.list | in.E5.list | in.E6.list | in.E7.list |
|---|---|---|---|---|---|---|---|
| chrX | ACE2 | NM_021804 | no | no | yes | no | no |
| chrX | AIFM1 | NM_001130847 | no | yes | yes | yes | yes |
| chrX | APEX2 | NM_001271748 | no | no | yes | no | no |
| chrX | APOO | NR_026545 | yes | yes | yes | no | no |
| chrX | ATP6AP1 | NM_001183 | no | yes | yes | yes | yes |
| chrX | BCAP31 | NM_001256447 | no | no | yes | no | no |
| chrX | BEX4 | NM_001080425 | no | yes | yes | yes | yes |
| chrX | BRWD3 | NM_153252 | no | yes | yes | no | no |
| chrX | CA5B | NM_007220 | no | yes | yes | no | no |
| chrX | CHM | NM_000390 | yes | no | yes | no | yes |
| chrX | COX7B | NM_001866 | no | no | yes | no | no |
| chrX | CXorf38 | NM_144970 | no | yes | yes | no | no |
| chrX | CXorf40B | NM_001013845 | no | no | yes | no | no |
| chrX | CXorf56 | NM_001170569 | yes | yes | yes | no | yes |
| chrX | DKC1 | NM_001142463 | yes | yes | yes | yes | no |
| chrX | DLG3 | NM_021120 | no | no | yes | no | no |
| chrX | EBP | NM_006579 | yes | yes | yes | yes | yes |
| chrX | EIF1AX | NM_001412 | no | yes | yes | yes | yes |
| chrX | EIF2S3 | NM_001415 | yes | yes | yes | yes | yes |
| chrX | EMD | NM_000117 | no | yes | yes | no | no |
| chrX | FAM127B | NM_001134321 | no | no | yes | no | yes |
| chrX | FMR1NB | NM_152578 | no | yes | yes | yes | yes |

| | | | | | | | |
|---|---|---|---|---|---|---|---|
| chrX | FTX | NR_028379 | no | no | yes | no | yes |
| chrX | FUNDC1 | NM_173794 | no | no | yes | yes | yes |
| chrX | FUNDC2 | NM_023934 | no | no | yes | yes | no |
| chrX | GAGE1 | NR_102272 | no | yes | yes | yes | yes |
| chrX | GAGE12J | NM_001098406 | no | yes | yes | yes | yes |
| chrX | GDPD2 | NM_001171192 | no | yes | yes | yes | no |
| chrX | HDAC6 | NM_006044 | no | no | yes | no | no |
| chrX | HDHD1 | NM_001135565 | no | no | yes | yes | yes |
| chrX | HMGB3 | NM_005342 | no | yes | yes | yes | yes |
| chrX | HNRNPH2 | NM_019597 | no | no | yes | no | no |
| chrX | HUWE1 | NM_031407 | no | no | yes | no | no |
| chrX | IDH3G | NM_174869 | no | no | yes | no | no |
| chrX | KIF4A | NM_012310 | no | no | yes | no | no |
| chrX | LAGE3 | NM_006014 | no | yes | yes | yes | yes |
| chrX | LAMP2 | NM_002294 | no | no | yes | yes | yes |
| chrX | MAGEA4 | NM_002362 | yes | yes | yes | yes | no |
| chrX | MAGEB2 | NM_002364 | no | yes | yes | yes | yes |
| chrX | MAGED2 | NM_177433 | no | no | yes | no | no |
| chrX | MAGT1 | NM_032121 | yes | no | yes | no | yes |
| chrX | MAP7D3 | NM_024597 | no | yes | yes | no | no |
| chrX | MCTS1 | NM_001137554 | no | no | yes | yes | yes |
| chrX | MECP2 | NM_001110792 | no | no | yes | no | no |
| chrX | MTRNR2L10 | NM_001190708 | yes | yes | yes | yes | yes |
| chrX | NONO | NM_001145410 | yes | yes | yes | yes | yes |
| chrX | NSDHL | NM_015922 | no | no | yes | yes | no |
| chrX | OFD1 | NM_003611 | no | no | yes | no | no |
| chrX | PDZD11 | NM_016484 | no | no | yes | no | no |
| chrX | PGK1 | NM_000291 | no | no | yes | yes | yes |
| chrX | PGRMC1 | NM_006667 | no | no | yes | no | yes |
| chrX | PLS3 | NM_005032 | no | yes | yes | yes | yes |
| chrX | PQBP1 | NM_001167992 | yes | yes | yes | no | no |
| chrX | PRKX | NM_005044 | no | no | yes | no | yes |
| chrX | PRPS2 | NM_002765 | no | no | yes | no | no |
| chrX | PSMD10 | NM_002814 | no | no | yes | no | no |
| chrX | RAB9A | NM_001195328 | no | yes | yes | no | no |
| chrX | RBM3 | NM_006743 | yes | yes | yes | yes | yes |
| chrX | RBMX | NM_001164803 | yes | yes | yes | yes | yes |
| chrX | RPL39 | NM_001000 | yes | yes | yes | yes | yes |
| chrX | SAT1 | NM_002970 | no | yes | yes | yes | yes |
| chrX | SLC25A5 | NM_001152 | yes | yes | yes | yes | yes |
| chrX | SLC25A53 | - | no | no | yes | no | no |
| chrX | SMC1A | NM_001281463 | no | no | yes | yes | yes |
| chrX | STS | NM_000351 | no | no | yes | yes | yes |
| chrX | TIMM17B | NM_005834 | no | yes | yes | yes | yes |
| chrX | TIMM8A | NM_001145951 | no | yes | yes | yes | yes |
| chrX | TKTL1 | NM_001145934 | no | yes | yes | no | no |
| chrX | TMSB4X | NM_021109 | no | yes | yes | yes | yes |
| chrX | TSIX | - | yes | yes | yes | yes | yes |
| chrX | TSPAN6 | NM_001278743 | no | yes | yes | yes | yes |
| chrX | UBL4A | NM_014235 | no | no | yes | yes | yes |
| chrX | VBP1 | NM_003372 | no | yes | yes | yes | yes |
| chrX | VCX | NM_013452 | yes | yes | yes | yes | yes |
| chrX | VCX2 | NM_016378 | yes | yes | yes | yes | yes |
| chrX | VCX3A | NM_016379 | yes | yes | yes | yes | yes |
| chrX | VCX3B | NM_001001888 | yes | yes | yes | yes | yes |
| chrX | WWC3 | NM_015691 | no | no | yes | no | no |
| chrX | XAGE3 | NM_133179 | no | yes | yes | yes | yes |
| chrX | XAGE5 | NM_130775 | no | no | yes | no | no |
| chrX | ZMAT1 | - | no | no | yes | no | no |
| chrX | ZNF280C | NM_017666 | no | no | yes | no | no |
| chrX | ZNF75D | NM_001185063 | no | no | yes | no | no |

**Genes called as biallelic at E6 from Moreira de Mello.**

| Chromosome | Gene symbol | RefSeq accession | in.E3.list | in.E4.list | in.E5.list | in.E6.list | in.E7.list |
|---|---|---|---|---|---|---|---|
| chrX | AIFM1 | NM_001130847 | no | yes | yes | yes | yes |
| chrX | ALG13 | NM_001039210 | no | no | no | yes | yes |
| chrX | ATP6AP1 | NM_001183 | no | yes | yes | yes | yes |
| chrX | BEX4 | NM_001080425 | no | yes | yes | yes | yes |
| chrX | CXorf65 | NR_033212 | no | no | no | yes | no |
| chrX | DKC1 | NM_001142463 | yes | yes | yes | yes | no |
| chrX | EBP | NM_006579 | yes | yes | yes | yes | yes |
| chrX | EIF1AX | NM_001412 | no | yes | yes | yes | yes |
| chrX | EIF2S3 | NM_001415 | yes | yes | yes | yes | yes |
| chrX | FMR1NB | NM_152578 | no | yes | yes | yes | yes |
| chrX | FUNDC1 | NM_173794 | no | no | yes | yes | yes |
| chrX | FUNDC2 | NM_023934 | no | no | yes | yes | no |
| chrX | GAGE1 | NR_102272 | no | yes | yes | yes | yes |
| chrX | GAGE12J | NM_001098406 | no | yes | yes | yes | yes |
| chrX | GDPD2 | NM_001171192 | no | yes | yes | yes | no |
| chrX | HDHD1 | NM_001135565 | no | no | yes | yes | yes |
| chrX | HMGB3 | NM_005342 | no | yes | yes | yes | yes |
| chrX | LAGE3 | NM_006014 | no | yes | yes | yes | yes |
| chrX | LAMP2 | NM_002294 | no | no | yes | yes | yes |
| chrX | MAGEA4 | NM_002362 | yes | yes | yes | yes | no |
| chrX | MAGEB2 | NM_002364 | no | yes | yes | yes | yes |
| chrX | MCTS1 | NM_001137554 | no | no | yes | yes | yes |
| chrX | MMGT1 | NM_173470 | no | yes | no | yes | no |
| chrX | MPP1 | NM_001166462 | no | no | no | yes | yes |
| chrX | MTRNR2L10 | NM_001190708 | yes | yes | yes | yes | yes |
| chrX | NONO | NM_001145410 | yes | yes | yes | yes | yes |
| chrX | NSDHL | NM_015922 | no | no | yes | yes | no |
| chrX | PDHA1 | NM_000284 | no | no | no | yes | yes |
| chrX | PGK1 | NM_000291 | no | no | yes | yes | yes |
| chrX | PLS3 | NM_005032 | no | yes | yes | yes | yes |
| chrX | PRPS1 | NM_001204402 | no | no | no | yes | yes |
| chrX | RBM3 | NM_006743 | yes | yes | yes | yes | yes |
| chrX | RBMX | NM_001164803 | yes | yes | yes | yes | yes |
| chrX | RPL39 | NM_001000 | yes | yes | yes | yes | yes |
| chrX | SAT1 | NM_002970 | no | yes | yes | yes | yes |
| chrX | SLC25A5 | NM_001152 | yes | yes | yes | yes | yes |
| chrX | SMC1A | NM_001281463 | no | no | yes | yes | yes |
| chrX | SMS | NM_004595 | no | no | no | yes | yes |
| chrX | STS | NM_000351 | no | no | yes | yes | yes |
| chrX | TIMM17B | NM_005834 | no | yes | yes | yes | yes |
| chrX | TIMM8A | NM_001145951 | no | yes | yes | yes | yes |
| chrX | TMSB4X | NM_021109 | no | yes | yes | yes | yes |
| chrX | TSIX | - | yes | yes | yes | yes | yes |
| chrX | TSPAN6 | NM_001278743 | no | yes | yes | yes | yes |
| chrX | UBA1 | NM_153280 | no | yes | no | yes | no |
| chrX | UBL4A | NM_014235 | no | no | yes | yes | yes |
| chrX | VBP1 | NM_003372 | no | yes | yes | yes | yes |
| chrX | VCX | NM_013452 | yes | yes | yes | yes | yes |
| chrX | VCX2 | NM_016378 | yes | yes | yes | yes | yes |
| chrX | VCX3A | NM_016379 | yes | yes | yes | yes | yes |
| chrX | VCX3B | NM_001001888 | yes | yes | yes | yes | yes |
| chrX | XAGE3 | NM_133179 | no | yes | yes | yes | yes |

**Genes called as biallelic at E7 from Moreira de Mello.**

| Chromosome | Gene symbol | RefSeq accession | in.E3.list | in.E4.list | in.E5.list | in.E6.list | in.E7.list |
|---|---|---|---|---|---|---|---|
| chrX | AIFM1 | NM_001130847 | no | yes | yes | yes | yes |
| chrX | ALG13 | NM_001039210 | no | no | no | yes | yes |
| chrX | ATP6AP1 | NM_001183 | no | yes | yes | yes | yes |
| chrX | ATP6AP2 | NM_005765 | yes | no | no | no | yes |
| chrX | BEX4 | NM_001080425 | no | yes | yes | yes | yes |
| chrX | CHM | NM_000390 | yes | no | yes | no | yes |
| chrX | CXorf56 | NM_001170569 | yes | yes | yes | no | yes |

| | | | | | | | |
|---|---|---|---|---|---|---|---|
| chrX | EBP | NM_006579 | yes | yes | yes | yes | yes |
| chrX | EIF1AX | NM_001412 | no | yes | yes | yes | yes |
| chrX | EIF2S3 | NM_001415 | yes | yes | yes | yes | yes |
| chrX | FAM104B | NM_138362 | no | no | no | no | yes |
| chrX | FAM127A | NM_001078171 | no | no | no | no | yes |
| chrX | FAM127B | NM_001134321 | no | no | yes | no | yes |
| chrX | FAM50A | NM_004699 | no | no | no | no | yes |
| chrX | FMR1NB | NM_152578 | no | yes | yes | yes | yes |
| chrX | FTX | NR_028379 | no | no | yes | no | yes |
| chrX | FUNDC1 | NM_173794 | no | no | yes | yes | yes |
| chrX | GAGE1 | NR_102272 | no | yes | yes | yes | yes |
| chrX | GAGE12J | NM_001098406 | no | yes | yes | yes | yes |
| chrX | HDHD1 | NM_001135565 | no | no | yes | yes | yes |
| chrX | HMGB3 | NM_005342 | no | yes | yes | yes | yes |
| chrX | LAGE3 | NM_006014 | no | yes | yes | yes | yes |
| chrX | LAMP2 | NM_002294 | no | no | yes | yes | yes |
| chrX | LDOC1 | NM_012317 | no | no | no | no | yes |
| chrX | MAGEB2 | NM_002364 | no | yes | yes | yes | yes |
| chrX | MAGEC1 | NM_005462 | no | no | no | no | yes |
| chrX | MAGED1 | NM_006986 | no | no | no | no | yes |
| chrX | MAGT1 | NM_032121 | yes | no | yes | no | yes |
| chrX | MCTS1 | NM_001137554 | no | no | yes | yes | yes |
| chrX | MID1 | NM_000381 | no | no | no | no | yes |
| chrX | MPP1 | NM_001166462 | no | no | no | yes | yes |
| chrX | MTRNR2L10 | NM_001190708 | yes | yes | yes | yes | yes |
| chrX | NONO | NM_001145410 | yes | yes | yes | yes | yes |
| chrX | PDHA1 | NM_000284 | no | no | no | yes | yes |
| chrX | PGK1 | NM_000291 | no | no | yes | yes | yes |
| chrX | PGRMC1 | NM_006667 | no | no | yes | no | yes |
| chrX | PIN4 | NM_001170747 | no | yes | no | no | yes |
| chrX | PIR | NM_003662 | no | no | no | no | yes |
| chrX | PLS3 | NM_005032 | no | yes | yes | yes | yes |
| chrX | PRKX | NM_005044 | no | no | yes | no | yes |
| chrX | PRPS1 | NM_001204402 | no | no | no | yes | yes |
| chrX | RBM3 | NM_006743 | yes | yes | yes | yes | yes |
| chrX | RBMX | NM_001164803 | yes | yes | yes | yes | yes |
| chrX | RPL39 | NM_001000 | yes | yes | yes | yes | yes |
| chrX | SAT1 | NM_002970 | no | yes | yes | yes | yes |
| chrX | SLC25A5 | NM_001152 | yes | yes | yes | yes | yes |
| chrX | SMC1A | NM_001281463 | no | no | yes | yes | yes |
| chrX | SMS | NM_004595 | no | no | no | yes | yes |
| chrX | STS | NM_000351 | no | no | yes | yes | yes |
| chrX | TIMM17B | NM_005834 | no | yes | yes | yes | yes |
| chrX | TIMM8A | NM_001145951 | no | yes | yes | yes | yes |
| chrX | TMSB4X | NM_021109 | no | yes | yes | yes | yes |
| chrX | TSIX | - | yes | yes | yes | yes | yes |
| chrX | TSPAN6 | NM_001278743 | no | yes | yes | yes | yes |
| chrX | TXLNG | NM_001168683 | no | no | no | no | yes |
| chrX | UBL4A | NM_014235 | no | no | yes | yes | yes |
| chrX | VBP1 | NM_003372 | no | yes | yes | yes | yes |
| chrX | VCX | NM_013452 | yes | yes | yes | yes | yes |
| chrX | VCX2 | NM_016378 | yes | yes | yes | yes | yes |
| chrX | VCX3A | NM_016379 | yes | yes | yes | yes | yes |
| chrX | VCX3B | NM_001001888 | yes | yes | yes | yes | yes |
| chrX | VGLL1 | NM_016267 | no | no | no | no | yes |
| chrX | XAGE3 | NM_133179 | no | yes | yes | yes | yes |
| chrX | ZFX | NM_003410 | no | yes | no | no | yes |
